# Visualizing ’omic feature rankings and log-ratios using Qurro

**DOI:** 10.1101/2019.12.17.880047

**Authors:** Marcus W. Fedarko, Cameron Martino, James T. Morton, Antonio González, Gibraan Rahman, Clarisse A. Marotz, Jeremiah J. Minich, Eric E. Allen, Rob Knight

## Abstract

Many tools for dealing with compositional “’omics” data produce feature-wise values that can be ranked in order to describe features’ associations with some sort of variation. These values include differentials (which describe features’ associations with specified covariates) and feature loadings (which describe features’ associations with variation along a given axis in a biplot). Although prior work has discussed the use of these “rankings” as a starting point for exploring the log-ratios of particularly high-or low-ranked features, such exploratory analyses have previously been done using custom code to visualize feature rankings and the log-ratios of interest. This approach is laborious, prone to errors, and raises questions about reproducibility. To address these problems we introduce Qurro, a tool that interactively visualizes a plot of feature rankings (a “rank plot”) alongside a plot of selected features’ log-ratios within samples (a “sample plot”). Qurro’s interface includes various controls that allow users to select features from along the rank plot to compute a log-ratio; this action updates both the rank plot (through highlighting selected features) and the sample plot (through displaying the current log-ratios of samples). Here we demonstrate how this unique interface helps users explore feature rankings and log-ratios simply and effectively.

## 1 Introduction

High-throughput sequencing and metabolomics data detailing the organisms, genes or molecules identified within a microbial sample are inherently compositional [1, 2]: that is, absolute abundances are often inaccessible and only relative information can be obtained from the data. These data must be interpreted accordingly. Performing a compositionally coherent analysis of how features change in a dataset generally requires selecting a “reference frame” (denominator) for log-ratio analysis, then relating the resulting log-ratios to sample metadata [2, 1].

Various tools for differential abundance analyses including but not limited to ALDEx2 [3] and Songbird [2] can produce *differentials*, which describe the log-fold change in relative abundance for features in a dataset with respect to certain covariate(s) [2]. Similarly, tools like DEICODE [4] can produce *feature loadings* that characterize features’ impacts in a compositional biplot [5]. Differentials and feature loadings alike can be sorted numerically and used as *feature rankings*, and this representation provides relative information about features’ associations with some sort of variation in a dataset [2, 4]. The natural next step is to use these rankings as a guide for log-ratio analyses (e.g. by examining the log-ratios of high-to low-ranked features). However, modern studies commonly describe hundreds or thousands of observed features: manually exploring feature rankings, whether as a tabular representation or as visualized using one-off scripts, is inconvenient.

Here we present Qurro (pronounced “churro”), a visualization tool that supports the analysis of feature log-ratios in the context of feature rankings and sample metadata. Qurro uses a two-plot interface: a “rank plot” shows how features are differentially ranked for a selected ranking column, and a “sample plot” shows log-ratios of the selected features across samples relative to selected sample metadata field(s). These plots are linked [6]: selecting features for a log-ratio highlights these features in the rank plot and updates the y-axis values of samples (corresponding to the value of the currently-selected log-ratio for each sample) in the sample plot. Due to its unique display, and the availability of multiple controls for feature selection and plot customization, Qurro simplifies the process of performing compositionally coherent analyses of ’omic data.

## 2 Implementation

Qurro’s source code is released under the BSD 3-clause license and is available at https://github.com/biocore/qurro.

Qurro’s codebase includes a Python 3 program that generates a visualization and the HTML/JavaScript/CSS code that manages this visualization. Qurro can be used as a standalone program or as a QIIME 2 plugin [7].

Both plots in a Qurro visualization are embedded as Vega-Lite JSON specifications [8], which are generated by Altair [9] in Qurro’s Python code. An advantage of Qurro’s use of the Vega infrastructure is that both plots in a Qurro visualization can be customized to the user’s liking in the Vega-Lite or Vega grammars. As an example of this customizability, the Vega-Lite specifications defining Figs. 1(a–c) and 2(a–c) of this paper were edited programmatically in order to increase font sizes, change the number of ticks shown, etc. (Our Python script that makes these modifications is available online; please see the “Data Availability” section.)

**Figure 1:**
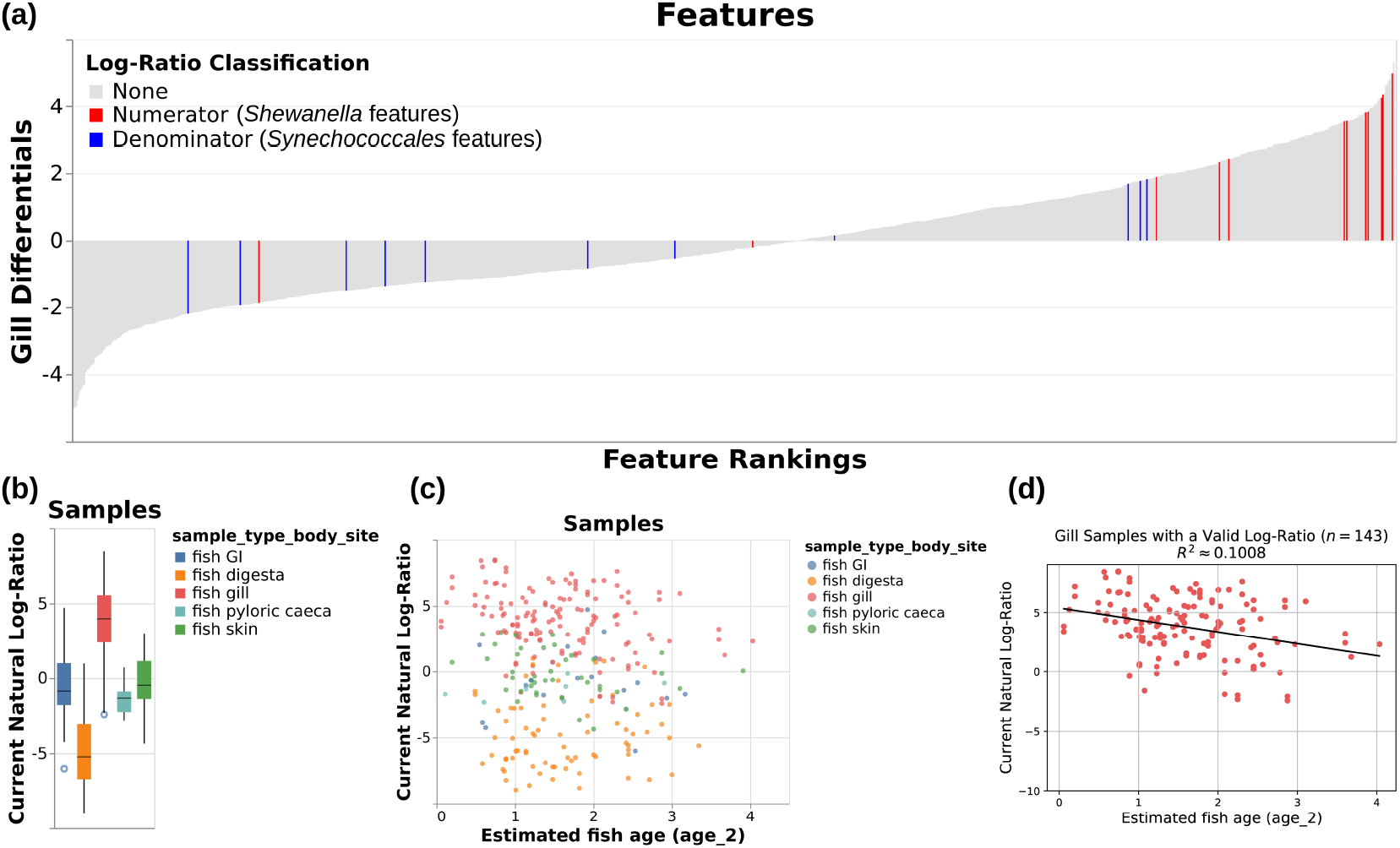
Various outputs from the case study showing the log-ratio of classified *Shewanella* features to classified *Synechococcales* features. **(a)** “Rank plot” showing differentials computed based on association with gill samples, using seawater samples as a reference. *Shewanella* features are colored in red, and *Synechococcales* features are colored in blue. **(b)** “Sample plot” in boxplot mode, showing samples’ *Shewanella-to-Synechococcales* log-ratios by sample body site. Note that only 285 samples are represented in this plot; other samples were either filtered out upstream in the analysis or contained zeroes on at least one side of their log-ratio. **(c)** “Sample plot,” showing a scatterplot of samples’ selected log-ratios versus estimated fish age. Individual samples are colored by body site. As in Fig. 1(b), only 285 samples are present. **(d)** Ordinary-least-squares linear regression (*R*^2^ ≈ 0.1008) between estimated fish age and the selected log-ratio for just the 143 gill samples shown in Fig. 1(b) and 1(c), computed outside of Qurro using scikit-learn [21] and pandas [12] and plotted using matplotlib [25].

**Figure 2:**
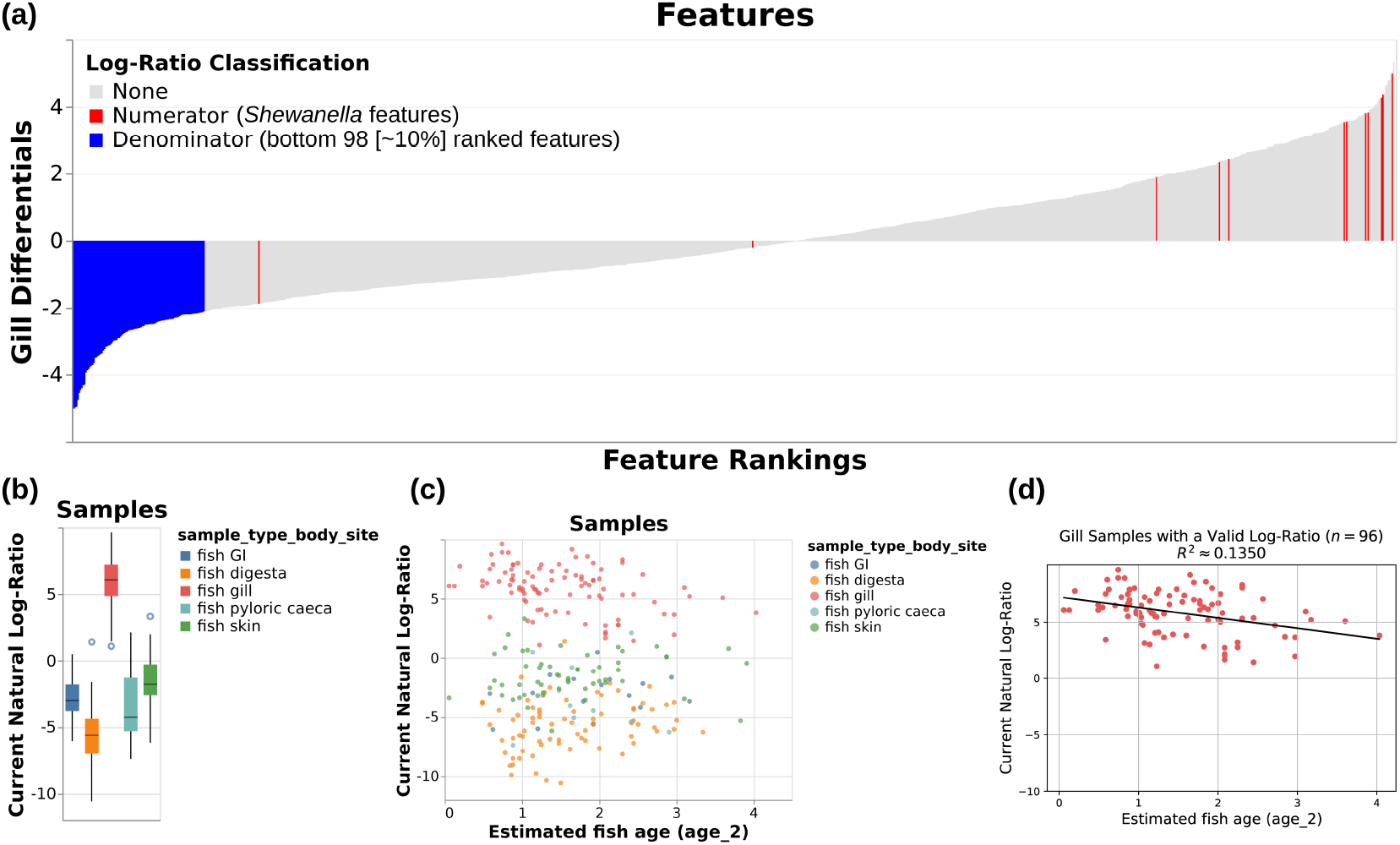
Various outputs from the case study (analogous to those in Fig. 1) showing the log-ratio of classified *Shewanella* features to the bottom 98 ranked features for the gill differentials. **(a)** “Rank plot” analogous to that shown in Fig. 1(a). **(b)** “Sample plot” in boxplot mode, showing the selected log-ratios of samples by body site. 252 samples are represented in this plot; as in Fig. 1(b), other samples were either filtered out upstream in the analysis or contained zeroes on at least one side of their log-ratio. **(c)** “Sample plot,” showing a scatterplot of samples’ selected log-ratios versus estimated fish age. Individual samples are colored by body site. As in Fig. 2(b), only 252 samples are present. **(d)** Ordinary-least-squares linear regression (*R*^2^ ≈ 0.1350) between estimated fish age and the selected log-ratio for just the 96 gill samples shown in Fig. 2(b) and 2(c), computed outside of Qurro as specified for Fig. 1(d).

### 2.1 Code dependencies

In addition to Altair, Qurro’s Python code directly relies on the BIOM format [10], Click (https://palletsprojects.com/p/click), NumPy [11], pandas [12], and scikit-bio (http://scikit-bio.org) libraries. Qurro’s web code relies on Vega [13], Vega-Lite [8], Vega-Embed (https://github.com/vega/vega-embed), RequireJS (https://requirejs.org), jQuery (https://jquery.com), DataTables (https://datatables.net), Bootstrap (https://getbootstrap.com), and Popper.js (https://popper.js.org).

## 3 Case study: the gills of *Scomber japonicus*

To demonstrate the utility of Qurro, we applied it to an extant dataset of V4-region 16S rRNA sequencing data from Pacific chub mackerel (*Scomber japonicus*) and environmental samples [14]. This dataset includes samples taken from five *Scomber japonicus* body sites (digesta, GI, gill, pyloric caeca, and skin) from 229 fish captured across 38 time points in 2017, along with many seawater, marine sediment, positive/negative control, and non-*Scomber japonicus* fish samples. A Jupyter Notebook [15] showing the steps taken during our re-analysis of this dataset is available online; see the “Data Availability” section.

### 3.1 Sample processing and re-analysis

When these samples were initially processed, the KatharoSeq protocol [16] was followed. This led us to exclude samples with less than 1,370 total counts from our re-analysis of this dataset.

Sequencing data (already processed using QIIME 1.9.1 [17] and Deblur [18] on Qiita [19]) were further processed and analyzed using QIIME 2 [7]. Taxonomic classification was performed using q2-feature-classifier’s `classify-sklearn` method [20, 21]. In particular, we trained a Naïve Bayes classifier on the SILVA 132 99% database [22] on sequences extracted using the same forward [23] and reverse [24] primers that were used for sample processing.

Due to upstream filtering in the re-analysis (a combination of filtering out non-*Scomber japonicus* and non-seawater samples, applying the aforementioned KatharoSeq sample exclusion criterion, Songbird’s default `--min-feature-count` of each feature needing to be present in at least 10 samples, and Qurro’s behavior of filtering out empty samples and features), 639 samples and 985 features were included in the Qurro visualization produced for this case study.

### 3.2 Computing “body site” differentials

One basic question about this dataset we decided to investigate using Qurro was of which features were associated with which *Scomber japonicus* body sites. To produce feature rankings accordingly, we used Songbird [2] to compute differentials detailing features’ associations with samples from each of the five studied body sites, using seawater samples in the dataset as a reference (supplementary information, section 1).

In general, highly-ranked features in a given differential column are positively associated with samples from that column’s corresponding body site, while lowly-ranked features are negatively associated with samples from that column’s corresponding body site (both in the context of the seawater samples). These differentials can be thought of as a starting point for investigating differentially abundant features for particular fish body sites in this dataset in a compositionally coherent manner.

### 3.3 Using Qurro to analyze differentials and log-ratios

Qurro simplifies the process of analyzing features’ differential abundances in the context of these differentials. The “rank plot” of a Qurro visualization is a bar plot where each bar corresponds to a single differentially ranked feature: the y-axis values of each bar are the literal differential or feature loading values, and features are sorted in ascending order by these values along the x-axis. The ranking column used is configurable, so Qurro users can quickly toggle among the input differentials or feature loadings; for the case study Qurro visualization, this means that users can—for example—switch between differentials computed based on association with skin samples to differentials computed based on association with gill samples.

#### 3.3.1 Highlighting features on the rank plot

The initial study of this dataset [14] agreed with prior work [26] on the frequency of *Shewanella* spp. in the fish gill microbiome. Qurro supports searching for features using arbitrary feature metadata (e.g. taxonomic annotations), and using this functionality to highlight *Shewanella* spp. on the rank plot of gill differentials (supplementary information, section 2) corroborates these findings: as Fig. 1(a) shows, the majority of identified *Shewanella* spp. are highly ranked in association with gill samples using seawater samples as a reference.

Particularly high-or low-ranked features like *Shewanella* spp. can merit further examination via a log-ratio analysis [2]; in particular, one question we might be interested in asking at this point is if *Shewanella* spp. are similarly abundant across other fish body sites. The remainder of this case study discusses a simple investigation in pursuit of an answer to this question, as well as to a few other questions that came up along the way.

#### 3.3.2 Choosing a suitable “reference frame.”

The compositional nature of marker gene sequencing data means that we cannot simply compare the abundances of *Shewanella* across samples in this dataset alone; however, we can instead compare the log-ratio of *Shewanella* and other features in this dataset across samples [2].

For demonstrative purposes, we chose the taxonomic order *Synechococcales* as the denominator (“reference frame”) for the first log-ratio shown here. Features in this dataset belonging to this order included sequence variants classified in the genera *Cyanobium*, *Prochlorococcus*, and *Synechococcus*. These are common genera of planktonic picocyanobacteria found ubiquitously in marine surface waters [27]. The expected stability of this group of features across samples in this dataset supports its use as a denominator here [2]. Furthermore, as shown in Qurro’s rank plot in Fig. 1(a), many *Synechococcales* features are relatively lowly ranked in association with gill samples (or at least relative to most *Shewanella* features); this gives additional reason to expect a comparative difference among gill samples for the *Shewanella*-to-*Synechococcales* log-ratio.

#### 3.3.3 Relating log-ratios to sample metadata

Upon selecting a numerator and a denominator for a log-ratio (in this case, by searching through taxonomic annotations), Qurro updates the sample plot so that all samples’ y-axis (“Current Natural Log-Ratio”) values are equal to the value of the selected log-ratio for that sample. The x-axis field, color field, and scale types of these fields—along with other options—can be adjusted by the user interactively to examine the selected log-ratio from a new perspective.

Once the log-ratio of *Shewanella*-to-*Synechococcales* was selected, Fig. 1(b) was produced by setting the sample plot x-axis to the categorical `sample_type_body_site` field and checking the “Use boxplots for categorical data?” checkbox. The resulting boxplot shows that the *Shewanella*-to-*Synechococcales* log-ratio is relatively high in gill samples, compared with other body sites’ samples (Fig. 1(b)). This observation corroborates the initial study of this dataset on the frequency of *Shewanella* particular to the fish gill microbiome [14].

Qurro can visualize quantitative sample metadata, as well. Using this functionality, we can add additional perspectives to our previously-reached observation. Age has been discussed as a factor impacting the microbiota of fish gills in this and other datasets [26, 14]. By setting the x-axis field to the `age_2` metadata column (estimated fish age), changing the x-axis field scale type to “Quantitative,” and setting the color field to `sample_type_body_site`, we get Fig. 1(c)—a scatterplot showing the selected log-ratio viewed across samples by the estimated age of their host fish.

One trend that stood out to us in this scatterplot was an apparent negative correlation between the selected log-ratio and estimated fish age for gill samples. To support further investigation of patterns like this, Qurro can export the data backing the sample plot to a standard tab-separated file format—this file can then be loaded and analyzed in essentially any modern statistics software or programming language. This functionality was used to generate Fig. 1(d), in which we quantify and visualize this correlation for gill samples using ordinary-least-squares linear regression (*R*^2^ ≈ 0.1008). Although obviously not evidence of a causal relationship, this result opens the door for further investigation of this trend. One of many possible explanations for this observed trend is that the gills of younger fish are differentially colonized by *Shewanella* spp. and/or by *Synechococcales*; this may, in turn, be reflective of factors like vertical habitat use, immune development, or food choice.

#### 3.3.4 Interrogating the “multiverse” of reference frames

Prior literature has shown the impact that choices in data processing can have on a study’s results, and on the corresponding “multiverse” of datasets generated during this process [28]. We submit that the choice of reference frame (denominator) in log-ratio analyses introduces a similar “multiverse”: for a set of *n* features, there are *O*(2*^n^*) possible subsets [2], so manually checking all possible reference frames for a given numerator is an intractable effort for the vast majority of datasets (although various heuristic methods have been proposed to address this sort of problem, e.g. [29]). In spite of this, the interactive nature of Qurro simplifies the task of validating results across reference frames.

Revisiting our analysis of *Shewanella* spp. in the gills of *Scomber japonicus*, there are multiple reasonable choices for reference frames. We chose *Synechococcales* mostly due to its expected ubiquity and stability across the marine samples in this dataset, but many other plausible choices exist.

In Fig. 2, *we repeat the exact same analysis as in Fig. 1*: but instead of using *Synechococcales* as the denominator of our log-ratio, we instead select the bottom ~ 10% (98/985) of features as ranked by gill differentials as the denominator (Fig. 2(a); supplementary information, section 2). Refreshingly, this log-ratio also shows clear “separation” of gill samples from other body sites’ samples in the dataset (Fig. 2(b)), as well as a similar negative correlation between estimated fish age and this log-ratio for gill samples (Figs. 2(c) and 2(d)) (*R*^2^ ≈ 0.1350). This serves as further evidence for our previous claims: although we still can’t say for sure, we can now more confidently state that *Shewanella* spp. seem to be dominant in the gills of *Scomber japonicus*, and that *Shewanella* abundance in these fishes’ gills seems to be negatively correlated with (estimated) fish age—since the trends shown in Figs. 1(b-d) and 2(b-d) have held up across multiple log-ratios with *Shewanella* as the numerator.

#### 3.3.5 Handling “invalid” samples

It is worth noting that many samples—including all of the seawater samples in the Qurro visualization (supplementary information, section 3)—are not present in Figs. 1(b–d) or 2(b–d). If a given sample in Qurro cannot be displayed for some reason—for example, the sample has a zero in the numerator and/or denominator of the currently selected log-ratio— Qurro will drop that particular sample from the sample plot. Furthermore, to make sure the user understands the situation, Qurro will update a text display below the plot that includes the number and percentage of samples excluded for each “reason.” This behavior helps users avoid spurious results caused by visualizing only a small proportion of a dataset’s samples.

### 3.4 Using Qurro in practice

Since Qurro visualizations are essentially just web pages it is trivial to host them online, thus making them viewable by anyone using a compatible web browser. As an example of this we have made Qurro visualizations of various datasets, including the case study’s, publicly available at https://biocore.github.io/qurro. We encourage users of Qurro to share their visualizations in this way, whenever possible, in order to encourage reproducibility and facilitate public validation of the conclusions drawn. Furthermore, we encourage readers of this paper to reconstruct Figs. 1 and 2 and verify that this paper’s claims are accurate.

## 4 Conclusions

Qurro serves as a natural “first step” for users of modern differential abundance tools to consult in order to analyze feature rankings, simplifying the work needed to go from hypothesis to testable result. We have already found it useful in a variety of contexts (e.g. [30]), and it is our hope that others find similar value.

As more techniques for differentially ranking features become available, we believe that Qurro will fit in as a useful piece within the puzzles represented by modern ’omic studies.

## 5 Data Availability

All data used was obtained from study ID 11721 on Qiita. Deblur output artifact ID 56427 was used, in particular. Sequencing data is also available at the ENA (study accession PRJEB27458). Various Jupyter Notebooks and files used in the creation of this paper are available at **https://github.com/knightlab-analyses/qurro-mackerel-analysis**.

## Acknowledgements

The authors thank Julia Gauglitz, Shi Huang, Franck Lejzerowicz, Robert Mills, Justin Shaffer, Seth Steichen, Bryn Taylor, and Yoshiki Vázquez-Baeza for helpful comments on the tool’s functionality and design. The authors thank Gail Ackermann for assistance with study metadata. The authors also thank Jake VanderPlas, Dominik Moritz, and the rest of the Altair/Vega development teams for answering questions regarding their software. Lastly, the authors thank Sarah Allard, Tomasz Kościółek, Franck Lejzerowicz, Anupriya Tripathi, and Yoshiki Vázquez-Baeza for help naming the tool.

## Funding

This work was supported in part by a University of California San Diego Computer Science and Engineering department fellowship to M.W.F.; the Joint University Microelectronics Program’s Center for Research on Intelligent Storage and Processing-in-memory, task number 2780.023, to M.W.F.; the IBM AI Horizons Network to M.W.F.; the University of California San Diego Frontiers of Innovation Scholars Program to C.M.; the National Science Foundation grant GRFP DGE-1144086 to J.T.M.; the National Institute of Dental and Craniofacial Research through F31 Fellowship 1F31DE028478 to C.A.M.; and a Center for Microbiome Innovation Microbial Sciences Graduate Research Fellowship to J.J.M.

## Conflict of interest statement

None declared.

## Notes

https://github.com/knightlab-analyses/qurro-mackerel-analysis

https://biocore.github.io/qurro

https://github.com/biocore/qurro

