## Supplementary Information for "Visualizing ’omic feature rankings and log-ratios using Qurro"

#### 1 1. Computing feature differentials using Songbird

As discussed in the main text, the initial focus of this re-analysis was on visualizing the associations of features with different *Scomber japonicus* body sites. To assess this, we ran Songbird [1] using the formula `C(sample_type_body_site, Treatment('sea water'))`. This produced six columns of differentials:

1. `Intercept`

2. `C(sample_type_body_site, Treatment('sea water'))[T.fish GI]`

3. `C(sample_type_body_site, Treatment('sea water'))[T.fish digesta]`

4. `C(sample_type_body_site, Treatment('sea water'))[T.fish gill]`

5. `C(sample_type_body_site, Treatment('sea water'))[T.fish pyloric caeca]`

6. `C(sample_type_body_site, Treatment('sea water'))[T.fish skin]`

The last five columns of differentials (2–6) describe the association of features with samples from each of the studied body sites, using seawater samples as a reference via Treatment coding.

(For reference, the fourth column, `C(sample_type_body_site, Treatment('sea water'))[T.fish gill]`, is what is shown in the rank plot sub-figures in the main text.)

The first column, `Intercept`, is less easily interpretable and not particularly relevant to the case study; this column is produced automatically by Patsy (**https://patsy.readthedocs.io**), the library used by Songbird to represent input formulae as design matrices.

Due to some current technical limitations, Qurro (as of writing) changes or removes certain special characters like `[` or `'` from field names. This is why the gill differential field name shown in Qurro—`C(sample_type_body_site, Treatment(sea water))(T:fish gill)`—has a slightly different name than it did in Songbird's output. (This behavior is documented in Qurro's README, which is distributed with its source code at **https://github.com/biocore/qurro**.)

### 2 2. Qurro log-ratio-selection controls used

The log-ratios selected in Figs. 1 and 2 were selected using Qurro's filtering controls in the following way.

#### 2.1 Fig. 1 (*Shewanella* to *Synechococcales*)

1. The numerator was selected by filtering to features where the `Taxon` field contained the text `Shewanella`.

2. The denominator was selected by filtering to features where the `Taxon` field contained the text `Synechococcales`.

##### 2.2 Fig. 2 (*Shewanella* to bottom $\sim 10\%$ features)

1. The numerator was selected by filtering to features where the `Taxon` field contained the text `Shewanella`.

2. The denominator was selected by filtering to features where the gill differential value—that is, `C(sample_type_body_site, Treatment(sea water))(T:fish gill)`—was less than `-2.102`. (This value was chosen in order to make the denominator include exactly the bottom 98 features.)

### 3 3. Details on Qurro (and Songbird) input data filtering

Running Qurro requires a few distinct input files (or QIIME 2 artifacts, if running it as a QIIME 2 plugin): a feature table, a "rankings" file, a sample metadata file, and optionally a feature metadata file.

If any features within the feature table are not present in the input rankings, then Qurro will not include these features in the output visualization (since they would not be displayable on the rank plot). This means that, although Qurro doesn't impose very strict filtering guidelines on its own by default, the filtering behaviors of upstream "ranking" tools will necessarily impact the amount of data shown in Qurro.

Since this impacts the case study, we go into detail about this behavior here.

##### 3.1 Songbird's `--min-feature-count`

For the case study dataset described in the manuscript, there were 23,253 features present in the feature table before running Songbird. However, Songbird applies a default `--min-feature-count` (i.e. the minimum number of samples a feature must appear in) of 10: this resulted in a large amount of features being removed from the visualization due to only appearing in a handful of samples. This is why there are just 985 features in the resulting Qurro visualization. (When generating a Qurro visualization, Qurro will output details explaining—if applicable—why certain samples/features have been removed from the visualization.)

#### 3.2 Why aren't there any seawater samples shown in the paper figures?

One of the things we noticed midway through this case study was that *Shewanella* spp., for the most part, did not appear in seawater samples. To help explain this, we prepared a Jupyter Notebook [2] that shows why these samples have been dropped. This notebook is available in the repository **https://github.com/knightlab-analyses/qurro-mackerel-analysis**.

##### 3.2.1 Non-numeric `age_2` values.

Since the `age_2` field refers to the estimated age of a sample's host fish, this field is not meaningful for non-fish samples like seawater. As shown in the notebook, all of the 50 seawater samples in our feature table have a non-numeric `age_2` value—this is one of the "reasons" Qurro has for dropping samples from the sample plot, and it explains why seawater samples cannot be shown in Figs. 1(c) or 2(c).

##### 3.2.2 Relative lack of *Shewanella* features.

As shown in the notebook, only one of the 50 seawater samples in our feature table included a feature classified as *Shewanella*. This particular *Shewanella* feature only appears in two samples in the feature table (including the aforementioned seawater sample), so it is not ranked by Songbird due to the default `--min-feature-count` described above.

From the Qurro visualization's perspective, then, none of the seawater samples contains any *Shewanella* features—so visualizing both of the log-ratios shown in the paper's case study will necessarily involve filtering out all of the seawater samples, unless imputation of some form were to be used. This is the reason why seawater samples are not shown in Figs. 1(b) and 2(b), and it's a reason (in addition to the `age_2` reason) why seawater samples are not shown in Figs. 1(c) and 2(c).

##### 3.2.3 Reflection on this phenomenon.

We note that the large amount of samples dropped here was likely caused in part by the nature of the case study. Since in general different body sites are expected to harbor different microbial communities, it makes sense that taxa common in one sample type might go almost or completely undetected in other sample types.

When looking for differentially abundant taxa across more subtly different sample categories (e.g. skin samples at different timepoints in the progression of atopic dermatitis, as shown in [1]), we expect that sample dropout like what we observed with seawater samples here will be less of an issue.
